# Theory of Cell Body Lensing and Phototaxis Sign Reversal in “Eyeless” Mutants of *Chlamydomonas*

**DOI:** 10.64898/2026.03.11.710819

**Authors:** Sumit Kumar Birwa, Ming Yang, Adriana I. Pesci, Raymond E. Goldstein

## Abstract

Phototaxis of many species of green algae relies upon directional sensitivity of their membrane-bound photoreceptors, which arises from the presence of a pigmented “eyespot” behind them that blocks light passing through the cell body from reaching the photoreceptor. A decade ago it was discovered that the spherical cell body of the alga *Chlamydomonas reinhardtii* acts as a lens to concentrate incoming light, and that in “eyeless” mutants of *Chlamydomonas* the consequence of that focused light reaching the photoreceptor from behind is a reversal in the sign of phototaxis relative to the wild type behavior. We present a quantitative theory of this sign reversal by completing a recent simplified analysis of lensing [Yang, et al., *Phys. Rev. E* **113**, 022401 (2026)] and incorporating it into an adaptive model for *Chlamydomonas* phototaxis. This model shows that phototactic dynamics in the presence of lensing is subtle because of the existence of internal light caustics when the cellular index of refraction exceeds that of water. During each period of cellular rotation about its body-fixed axis, the photoreceptor receives two competing signals: a relatively long, slowly-varying signal from the direct illumination, and a stronger, shorter, rapidly-varying lensed signal. The reversal of the sign of phototaxis is then a consequence of the dominance of the flagellar photoresponse to the signal with the higher time derivative. These features lead to a quantitative understanding of phototaxis sign reversal, including bistability in the direction choice, a prediction that can be tested in single-cell tracking studies of mutant phototaxis.

## I. INTRODUCTION

Phototaxis exhibited by many species of green algae serves as a paradigm for the general problem of directional sensing by simple uni- and multicellular organisms [1]. Unlike the gradient climbing dynamics found, for example, in bacterial chemotaxis [2], motion of aneural protists in response to light signals is generally understood to be based on a “line-of-sight” mechanism in which light falling on a membrane-bound photoreceptor just below the outer cell wall triggers changes in the beating dynamics of flagella that turn the cell toward the light [3]. Phototaxis can be of either sign, depending on light levels and the presence of various chemical species in the surrounding medium [4]. For biflagellated unicellular organisms such as *Chlamydomonas reinhardtii* (CR), the two flagella respond in opposite ways to the internal chemical changes triggered by varying illumination of the photoreceptor [5]. As evidenced by the phototactic inability of the mutant *ptx1* which has symmetrical flagellar responses to light [6], this asymmetry is required for accurate phototaxis.

A long series of studies [3, 5, 7–9] has firmly established that CR phototaxis along with that of its multicellular relatives in the volvocine lineage—the 16-cell *Gonium* [10] and the 1000 − 2000 cell *Volvox* [11]—can be understood quantitatively as a consequence of the interplay of two universal features: persistent rotational motion of the organisms about a body-fixed axis and the directionally sensitivity of the photoreceptor. The spinning motion arises from particular broken symmetries in the beat pattern of the flagella, for example a slight out-of-plane breaststroke beating of the CR flagella [12] or a tilt of the beat plane relative to the anterior-posterior axis of *Volvox*. The directionality of the photoreception arises from the presence of a pigmented “eyespot” behind the photoreceptor, a structure composed of carotenoid globules [13] that is thought to act like a quarter-wave plate due to its thickness relative to the wavelength of light [14] so as to reflect light (by constructive interference) as shown in Fig. 1. In this view, light incident on the front of the cell from the surrounding water would pass once through the photoreceptor, be reflected by the eyespot and pass through the photoreceptor a second time, increasing the signal in a manner analogous to the way in which the *tapetum lucidum* behind the retina of animals such as dogs, cats and fish [15] retroreflects light. Likewise, light passing through the cell body from behind is blocked from reaching the receptor by the same reflective action of the eyespot.

**FIG. 1.**
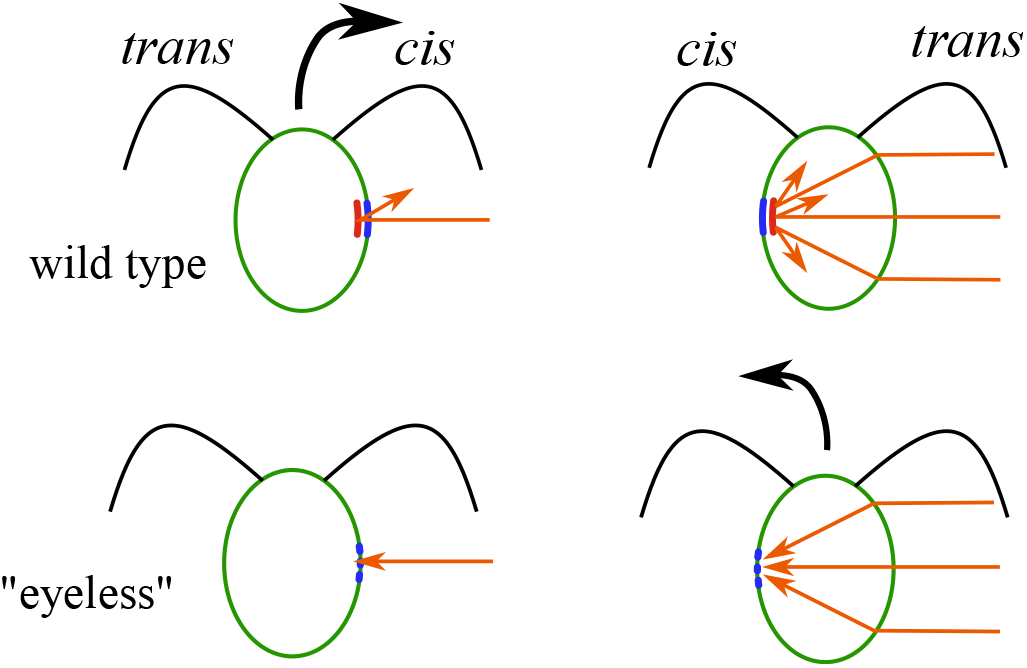
Phototaxis of *Chlamydomonas*. Photoreceptor is shown in blue, eyespot in red. Adapted from [19] and [20].

The role of the eyespot in shielding the photoreceptor has been the subject of debate, with some evidence [16] indicating a rather narrow wavelength-dependent peak in reflectivity, favoring the alternate possibility that receptor directionality arises from the absorptive properties of chlorophyll within the cell body. An earlier study [17] of mutants lacking chlorophyll and the eyespot investigated the effect of reconstituting photoreceptors with externally added retinal. Strikingly, the sign of phototaxis observed was opposite to that of the wild type, a result suggesting that lensing by the cell body might concentrate light incident from behind the cell.

Further evidence for the possibility of cell body lensing was presented in the important work by Kessler, et al. [18], followed by the key work of Ueki, et al. [19] that not only demonstrated the focusing effect directly, with an estimate of ≈1.47 for the effective index of refraction *n*_*c*_ of the cell body, but also showed that a number of eyeless mutants had reversed-sign phototaxis relative to the wild type. These results strongly support the role of the eyespot in blocking light from behind the photoreceptor.

Motivated by cell body lensing effects in simple geometries, recent work [20] has considered the general problem of light refraction by complex shapes found in the world of protists. It also used geometric optics to estimate the intensity boost *η* associated with lensing of light, defined as the ratio of light falling on the photoreceptor due to lensing relative to that incident from in front of the cell. For the special case of light coming from directly behind a spherical cell, a considerable boost of *η* ≈ 1.5 was found.

It follows from this result that as a swimming eyeless mutant spins around its axis, instead of just receiving the single pulse-like half-wave-rectified-sinusoid signal of the wild type, it receives a second, shorter but more intense signal during each full rotation. It is thus not at all clear how eyeless cells make a decision as to which signal to follow. Do they simply follow the instantaneously stronger signal? Is there a threshold of amplification due to lensing for the opposite sign of phototaxis to occur? How precise is the sign-reversed phototaxis? These questions are examples of the more general issue of decision-making by aneural organisms confronted by competing stimuli, and have been recently studied in detail for CR responding to two separate beams of light with adjustable amplitude and angular separation [21].

In this paper we present a theoretical analysis of the phototactic response of eyeless mutants by generalizing the earlier optical analysis [20] mentioned above to include off-axis lensing effects and then incorporating those results into the adaptive model [9] of CR phototaxis. The lensing of incoming parallel light within a spherical refractive cell consists of an illuminated cone, outside of which is dark (see Fig. 3(b) below). The outer cone boundary is a caustic, with a well-known peak in intensity. This light pattern within the cell must be taken into account to determine the phototactic steering of a cell. We show that even in the simplest treatment of these features there is competition in phototactic direction due to the two signals; cells can exhibit wild-type or sign-reversed phototaxis depending on their initial orientation with respect to the direction of the incident light.

## II. LENSING

A quantitative theory of phototaxis requires knowledge of the light intensity falling on the photoreceptor as a function of incident angle. The photoreceptor occupies a roughly circular patch of the cell membrane, with a radius *a* ∼ 0.4 − 0.7 *µ*m [14, 22] as shown in Fig. 1 that is considerably smaller than the cell radius *R* ≈ 5 *µ*m, and thus the receptor may be considered a planar disk. Furthermore, the smallness of the ratio

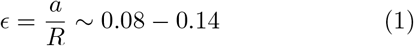

justifies the limit *ϵ* → 0 adopted in calculations below.

We assume the cell is spherical and has a uniform index of refraction *n*_c_ ≥ *n*_*w*_, where *n*_w_ ≃ 1.33 is the index of refraction of the surrounding aqueous medium. The experimental results of Ueki, et al. [19] suggested that the internal index is *n*_c_ ≃ 1.47, which indicates the relative refractive index *n* = *n*_c_*/n*_w_ ≃ 1.1, a representative value we use in numerical studies of phototaxis.

Consider the situation in which a cell swims toward +*y* in the *xy*-plane while light of intensity *I*_0_ shines toward +*x* as in Figure 2, where the cell is viewed from behind and rotates counterclockwise. Choose the origin of coordinates to be the center *O* of the cell and let *ζ* ∈ [0, 2*π*] be the angle of rotation around about *y* (which is coincident with the direction ê _3_ of the body-fixed posterior-anterior axis), whose origin is chosen such that the angle indicating the location of the eyespot midpoint is also *ζ*.

**FIG. 2.**
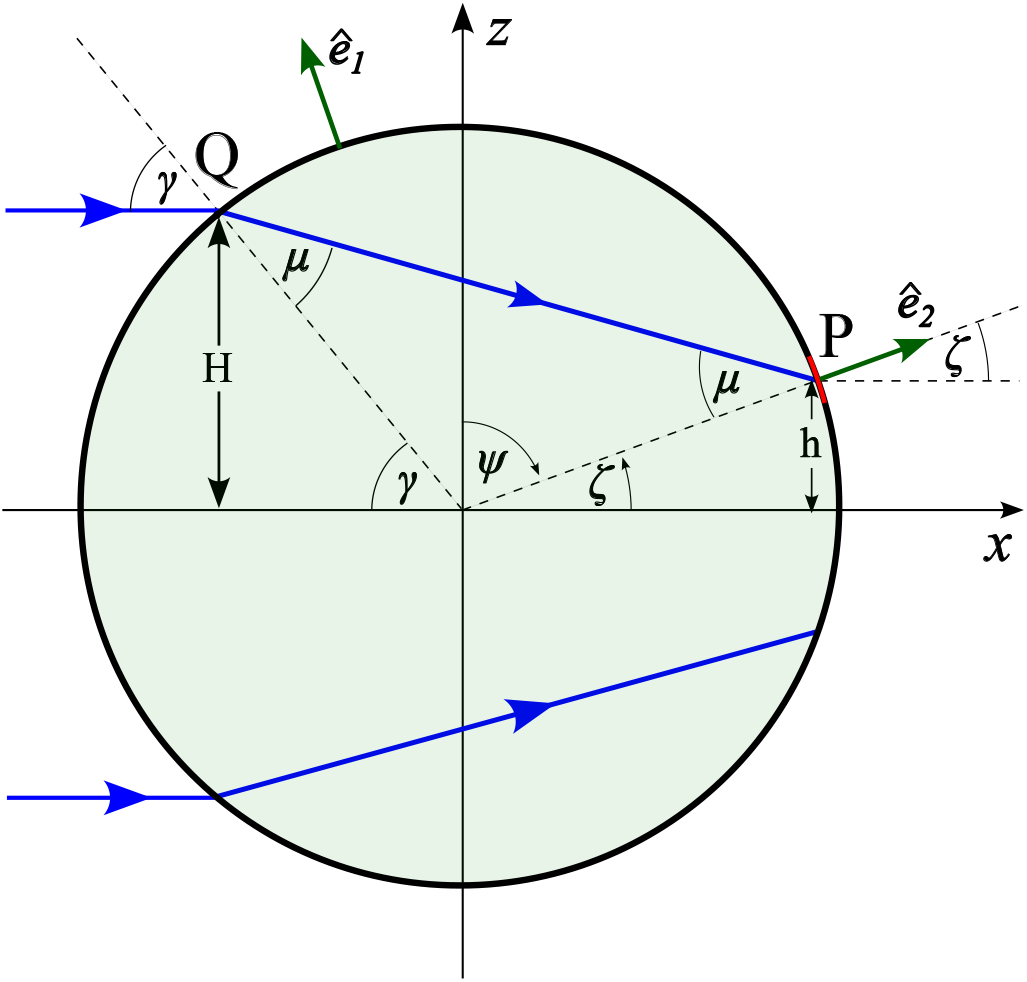
Cross section of a spherical cell illuminated from the left. Cell swims with its body-fixed axis **ê**_3_ along the positive *y*-axis and rotates counterclockwise when viewed from behind, as in the drawing. Visible body-fixed axes are **ê**_1_ and **ê**_2_. The top ray of light (blue) enters the cell at *Q* and intercepts the photoreceptor (red arc) at *P* .

For angles in the range *π*/2 ≤ | *ζ* | ≤ *π* light is incident on the photoreceptor from the outside, with an average projection *I*(*ζ*) = |cos *ζ*|, reaching a maximum when the midpoint of the photoreceptor is on the *x* axis.

The quantity of interest now is the intensity *I*(*ζ*) for all other possible values of *ζ* when the light impinges on the photoreceptor from behind, having passed through the cell body. In previous work [20], we studied the high-symmetry case in which the photoreceptor midpoint is again on the *x* axis, but on the far side of the cell (*ζ* = 0). In the limit *ϵ* ≪ 1, neglecting contributions from internal reflections, the on-axis intensity boost *η*_0_ ≡ *I*(0) is [20]

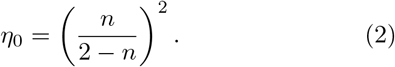

As mentioned above, for CR with *n* = 1.1, *η*_0_ has the surprisingly large value ∼ 1.5. (Note: *n* = 2 corresponds to the lowest value at which the focal point of paraxial rays is inside the sphere.)

To find the intensity boost for all values of *ζ*, we must find (see Fig. 2) which one of the incoming parallel lights rays refracts at point *Q* on the cell boundary and impinges on the photoreceptor at *P* . The angles of incidence and refraction with respect to the surface normal at *Q* are *γ* and *µ*, respectively, and they obey Snell’s law,

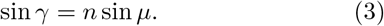

From elementary geometry we find

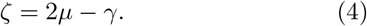

Viewing *ζ* as the independent variable, Eqs. (3) and (4) provide two equations for the two unknowns *γ*(*ζ*) and *µ*(*ζ*) which are solved by the implicit equations

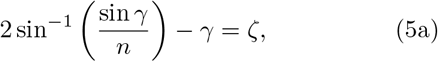

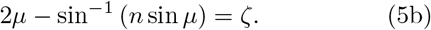

For *n* = 1, when there is no lensing, *γ* = *µ* = *ζ*, as expected. For small *ζ*, one finds the expansions

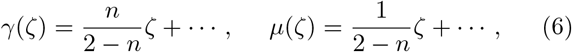

which are useful in determining the on-axis limit of the more general lensing expression below.

Figure 3(a) shows the function *γ*(*ζ*) for *n* = 1.1, and illustrates that there is a maximum possible value *ζ*_*m*_ for *γ* ∈ [0, *π/*2]. The physical significance of this is seen in Fig. 3(b), which displays ray-tracings [23] for a circular lens with *n* = 1.1. We see that the maximum *ζ* is associated with the formation of caustics; no light reaches the photoreceptor for *ζ*_*m*_ *<* |*ζ*| *< π/*2. We refer to this annular domain as the “dark region”.

**FIG. 3.**
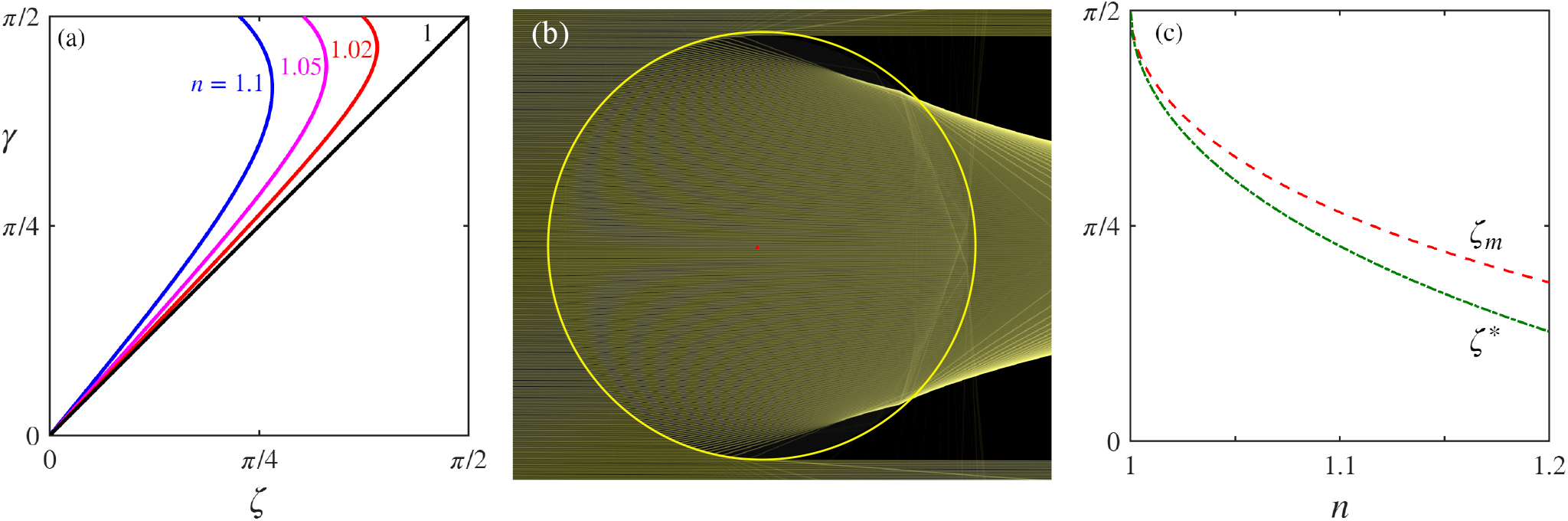
Geometrical optics for spherical cells. (a) Relationship (5) between the incident angle *γ* and the angle *ζ* of the photoreceptor shown in Fig. 2. (b) Ray-tracings [23] illustrating caustic formation for relative index of refraction *n* = 1.1. (c) The angle *ζ*_*m*_ of the caustic and angle *ζ*\* bounding the double-valued region of light, as function of the relative index *n*.

The maximum angle *γ*_*m*_ associated with *ζ*_*m*_ as a function of index of refraction ratio *n* can be found by the condition *dζ/dγ* = 0, where

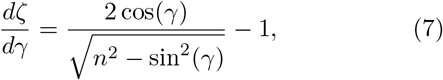

yielding

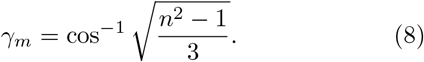

A second characteristic angle is associated with an incident beam that grazes the top of the circle, with incident angle *γ*\* = *π/*2, corresponding to

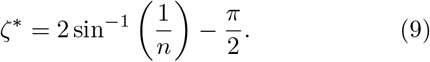

For photoreceptor angles | *ζ* |*< ζ*\* there is a unique incident ray that impinges on the photoreceptor, while for *ζ*\* *<*| *ζ* |*< ζ*_*m*_, two distinct incident light rays, with two distinct angles *γ*, refract to the same point given by *ζ*. As an example, for *n* = 1.1 we find *γ*_*m*_ ≈ 74.5°, *ζ*_*m*_ ≈ 47.8° and *ζ*\* ≈ 40.8°. The dependence of *ζ*_*m*_ and *ζ*\* on *n* is shown in Fig. 3(c).

With these results, we may now compute the intensity boost for a vanishingly small photoreceptor whose midpoint is at angle *ζ*. We do this by calculating the ratio between the size of the incoming bundle of light rays that ultimately impinges on the photoreceptor to the size of the receptor itself. From Fig. 2, let *H* = *R* sin *γ* be the the distance from the *x*-axis to *Q* and *h* = *R* sin *ζ* be the distance from the *x*-axis to *P* . Under a small change *δγ*, viewing *ζ* as a function of *γ*, we find

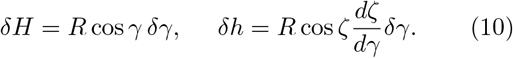

From trigonometry, we find that the bundle of rays heading from *Q* to *P* has a cross-sectional width *δh*_⊥_ = *δh* cos *µ/* cos *ζ*, from which it follows that

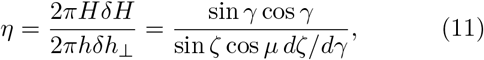

and the projection *I* of the refracted beam onto the photoreceptor is simply *I* = *η* cos *µ* or

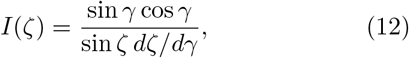

where *γ*(*ζ*) is the solution to the implicit equation (5a). The result (12) applies directly for |*ζ*| ∈ [0, *ζ*\*] where the function *γ*(*ζ*) is single-valued. In the interval |*ζ*| ∈ [*ζ*\*, *ζ*_m_] one must add the separate contributions from the two branches of *γ*(*ζ*).

Figure 4 shows the complete angular dependence of the light intensity, both for illumination and the lensed regions, for several values of *n*. For any *n >* 1 we see that the intensity diverges at the angle *ζ*_*m*_, the location of the caustic, because the derivative *dζ/dγ* = 0 there. But around *ζ* = 0 the lensed intensity is just a magnified form of the underlying cos *ζ* dependence; the dashed blue line in Fig. 4 is the function *η*_0_ cos *ζ*, with *η*_0_ given by (2). Additionally, one verifies from (6) and (12) that the on-axis boost result (2) is recovered as *ζ* → 0, as when *γ, µ, ζ* ≪ 1, sin *γ/* sin *ζ* ≈ *γ/ζ* ≈ *dγ/dζ* = *n/*(2 − *n*), and cos *γ* ≈ 1, so *I*(0) = (*n/*(2 − *n*))^2^.

**FIG. 4.**
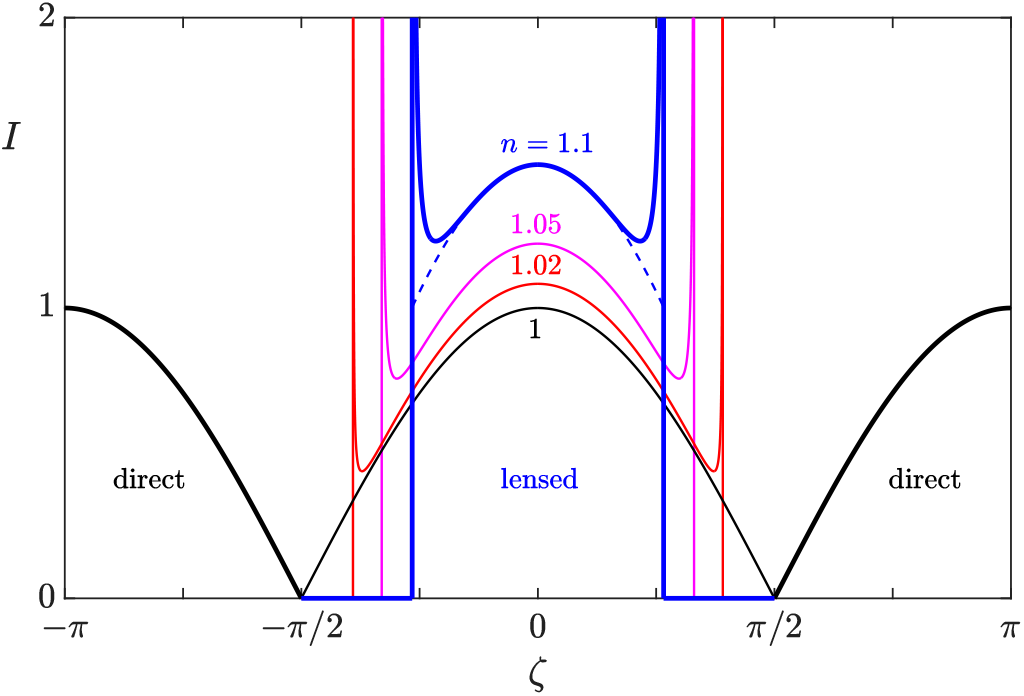
Angular dependence of lensing. Intensity versus rotation angle *ζ* for various values of *n*. Direct contributions are shown as heavy black curves. Dashed blue line is local approximation near *ζ* = 0 for representative value *n* = 1.1.

The cylindrical symmetry of the setup shown in Fig. 2 implies that pattern of lensed light within the cell is axisymmetric about *x*. In §III we show how this symmetry can be used to find the light intensity at the photoreceptor when it is out of the *xz*-plane.

## III. ADAPTIVE PHOTOTAXIS WITH LENSING

We now embed the calculation of the previous section in a theory of *Chlamydomonas* phototaxis [9], illustrating the result in the context of the simplest geometry of a phototurn: cells swimming in the *xy*-plane with a collimated source of light shining in the positive *x* direction.

In the model, a full account of swimming motions involves coupling the general rigid-body equations of motion for the three Euler angles (*ϕ, θ, ψ*) with dynamics describing how the relevant angular velocities (*ω*_1_, *ω*_2_, *ω*_3_) about the body-fixed axes (**ê**_1_, **ê**_2_, **ê**_3_) evolve as light falls on the photoreceptor and the flagellar beating dynamics change accordingly. With reference to Fig. 5, we choose **ê**_1_ to point normal to the plane containing the two flagella, **ê**_2_ to point towards the *cis* flagellum (that which is closer to the eyespot/photoreceptor) and **ê**_3_ to point normal to the cell anterior, in the direction of swimming. The eyespot is generally located close to the cell midplane, at some angle *κ* with respect to **ê**_2_ so that the eyespot normal is **ô** = sin *κ* **ê**_1_ + cos *κ* **ê**_2_ [9]. The results we obtain below are insensitive to the value of *κ* and for concreteness we set *κ* = 0, so **ô** = **ê**_2_, as in Fig. 2.

**FIG. 5.**
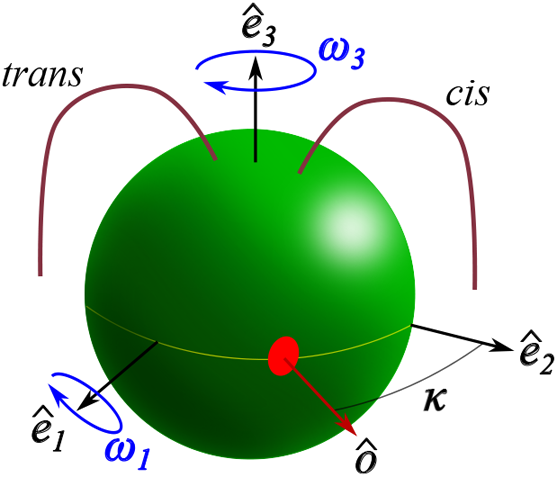
Coordinate system of *Chlamydomonas*, adapted from [21]. Eyespot is shown as red disc, with outward normal **ô**. Cell rotates continuously around **ê**_3_ and transiently around **ê**_1_ due to the photoresponse of its *cis* and *trans* flagella.

Because of the planarity of the flagellar beating, we may neglect any rotations around **ê**_2_, thus setting *ω*_2_ = 0 and focus only on (i) steady spinning of the cell around **ê**_**3**_ at constant frequency *ω*_3_ *<* 0 and (ii) the changes to *ω*_1_ arising from differences in the waveforms of the *cis* and *trans* flagella induced by changing illumination of the photoreceptor. Ignoring small out-of-plane motions, we fix *θ* = *π/*2 [9], and thus specialize to motions solely in the *xy*-plane, and find that the equations of motion for the Euler angles reduce to

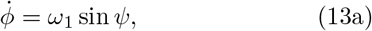

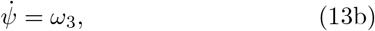

where *ϕ* is the angle between **ê**_3_ and the negative *y*-axis and *ψ* is the angle between **ê**_2_ and the *z*-axis. The adaptive dynamics is

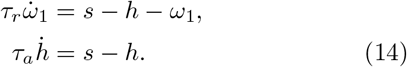

Here, *h* is a hidden variable that accounts for adaptation, *τ*_*r*_ and *τ*_*a*_ are the (short) response and (long) adaptation time scales governing beating asymmetries, and 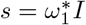 is the signal, governed by the light intensity *I* at the photoreceptor, parameterized in magnitude by the maximum angular frequency 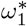 induced by the direct illumination.

We adopt the rescalings *T* = |*ω*_3_| *t, P* = *ω*_1_*/* |*ω*_3_|, 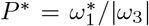, *α* = |*ω*_3_| *τ*_*a*_, *β* = |*ω*_3_| *τ*_*r*_, *H* = *h/* |*ω*_3_| and *S* = *s/* |*ω*_3_| . Integrating *ψ*_*T*_ = −1 and considering the relationship *ζ* = *π/*2 − *ψ* between the Euler angle *ψ* and the angle *ζ* in Fig. 2, we choose an integration constant in (13b) for later convenience and obtain *ζ*(*T* ) = *T* . As in previous work [21], we set *ϕ* = *π/*2 + *φ*, where *φ* is the angle between the swimming direction and the positive *x*-axis. The dynamics becomes

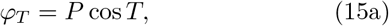

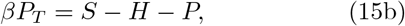

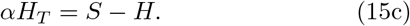

When the light shines toward the positive *x*-axis 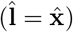, and the normal vector to the photoreceptor is

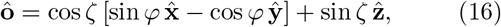

the signal *S* is related to the projection

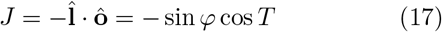

through

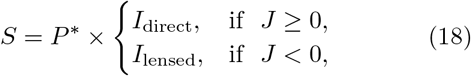

where *I*_direct_ = *J* and *I*_lensed_ is the result for arbitrary *φ* of the calculation presented in §II for the special case *φ* = *π/*2. To find the general expression, note that the axisymmetry of the lensed light implies that the intensity depends only on the angle *ζ*_eff_ between the eyespot normal and the positive *x*-axis. Simple geometry shows that

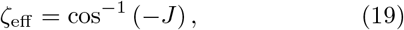

where *J* is given by (17). It follows that *I*_lensed_ is given by (12) with the substitutions *ζ* → *ζ*_eff_ and *γ* → *γ*_eff_, where *ζ*_eff_ and *γ*_eff_ also satisfy (5a). In the limit *n* → 1, where the lensing effects vanish and the light from behind the cell goes straight through, we have *ζ*_eff_ = *γ*_eff_ and thus *I* → cos *ζ*_eff_ = sin *φ* cos *T*, as expected.

In this rescaled system of units the swimming trajectories **R**(*T* ) are obtained by integrating the equations

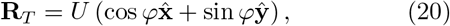

where *U* = *u/*(*R*| *ω*_3_| ) is the swimming speed *u* scaled by the cell radius *R* and rotational frequency. The model is now completely defined by Eqs. (15), (17)-(20), with parameters *α, β, P* *, *n*, and *U* .

We now turn to numerical results on phototaxis in the presence of lensing, show that lensing indeed produces a reversal in the sign of phototaxis, and propose a physical explanation for the effect. We continue with the geometry assumed in the previous section: a cell swimming in the *xy*-plane with light shining along the positive *x*-axis. Figure 6 shows (dark blue) the trajectory of a cell which starts at the origin with an initial orientation angle *φ* = 0.55 × *π*, so that it faces slightly toward the light. In Fig. 6(a) we consider the wild-type cell with eyespot shading and hence no lensing. The cell rapidly turns toward the light and ultimately swims toward it directly along the negative *x*-axis (*φ* = *π*). Figure 6(b) shows the projected angle *ζ*_eff_ versus time, illustrating that during the first few cellular rotations some lensed light would have been experienced were it not for eyespot shading, but eventually *ζ*_eff_ → *π/*2 as the cell aligns with the light and the eyespot is orthogonal to the light. The gray trajectories shown in Fig. 6(a), for initial angles uniformly spaced within the interval [0, *π*], illustrate that the positive phototaxis holds for all initial conditions.

**FIG. 6.**
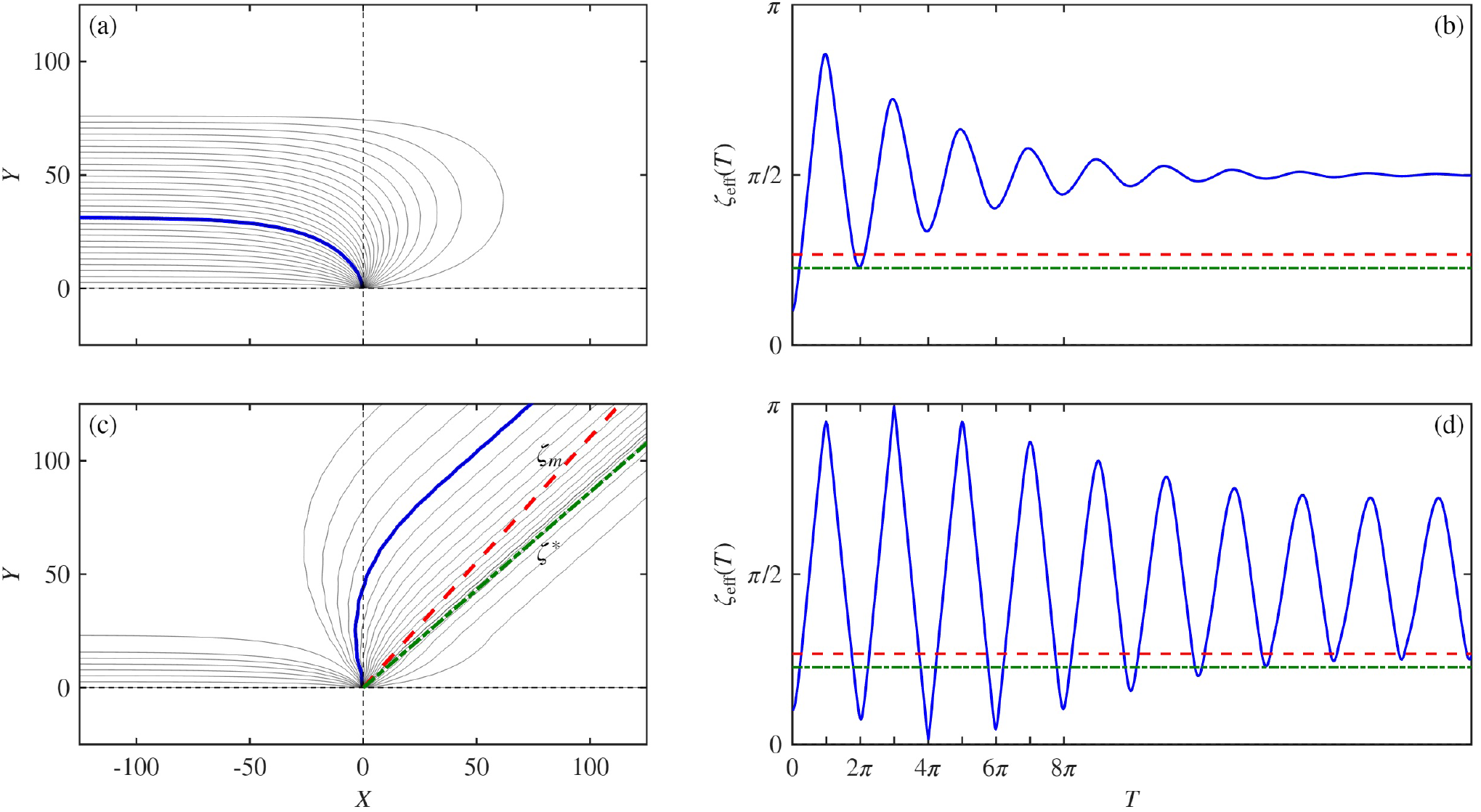
Numerical results. (a) Trajectories in *XY* -plane for a range of initial orientations *φ*∈ [0, *π*] for a positively phototactic cell without lensing, with parameters *α* = 2, *β* = 0.5, *P* * = − 0.4 and *U* = 2. Trajectory for initial condition *φ*(0) = 0.55*π* is shown in blue and the associated projection angle *ζ*_eff_ is shown in (b), where dashed lines indicate angles *ζ*_*m*_ and *ζ*\*. All initial orientations lead to positive phototaxis. (c) Trajectories as in (a) but with lensing. A small subset of initial orientations lead to positive phototaxis, but the majority produce negative phototaxis. Blue trajectory in (c) has same initial orientation as in (a). (d) As in (b), but with lensing. Dashed lines in (c) are lines at angles *ζ*_*m*_ and *ζ*\*.

In contrast to the wild-type behavior, Fig. 6(c) shows the trajectory (dark blue) for the same initial condition when lensed light is sensed. Although initially heading slightly towards the light, the cell turns around and swims away from it, exhibiting negative phototaxis and settling into a straight trajectory whose angle *φ*\* is between the two angles *ζ*\* and *ζ*_*m*_ associated with the lensing effect. This is seen clearly in Fig. 6(d), where, during each rotation of the cell, *ζ*_eff_ reaches a minimum between *ζ*\* and *ζ*_*m*_ and a maximum that is well beyond *π/*2. Thus, in the steady state swimming at long times, the photosensor detects two signals during each cellular rotation. These two signals induce phototaxis in opposite directions and those two effects balance to yield the compromise direction *φ*\*, whose precise value depends on the position of the photoreceptor relative to the cell midplane [21] and the relative index *n*. As in Fig. 6(a), trajectories for a range of initial orientations are shown in gray, and indicate that there is a small range of initial angles near *π* which lead to positively phototactic swimming. The great majority of initial conditions leads to negative phototaxis.

In order to understand these results, we note that the chief difference between the incident and lensed profiles is that the latter is confined to a smaller angular domain (see Figs. 3 and 4) and not only has a larger amplitude within that domain, but also has much higher derivatives due to the presence of the caustic at the domain edges. This difference holds in spite of the fact that the integrated flux of light incident on the cell is identical to the integrated lensed flux incident on the interior wall of the cell. We thus conclude that the reverse in sign of phototaxis is the consequence of the dominance of the lensed photoresponse over the direct one due to the larger time derivative of the lensed signal, as explained below.

That it is the time derivative of the signal that determines the flagellar photoresponse can be traced back to the pioneering work of Rüffer and Nultsch [5] who found that the *cis* flagellum increases (decreases) its beating amplitude during a step up (down) in light intensity, while the *trans* flagellum does the opposite. This asymmetry is necessary for wild-type phototaxis, as evidenced by the mutant strain *ptx1* which lacks this asymmetry and does not do proper phototaxis [6]. The role of this asymmetry in phototaxis can be seen by considering the photoresponse as the photoreceptor rotates first toward the light and then away, so the derivative *S*_*T*_ changes sign. The changes in flagellar beating when *S*_*T*_ *>* 0 must be opposite those when *S*_*T*_ *<* 0 half a turn later, or else the two responses will cancel and there will be no net turn toward the light.

The observation of Rüffer and Nultsch is actually embodied in the adaptive model of flagellar dynamics, even though the dynamics in (15b) and (15c) are forced by the signal *S* itself. As shown elsewhere [21], these coupled equations for *P* and *H* can be recast as a single equation for the photoresponse variable *P*,

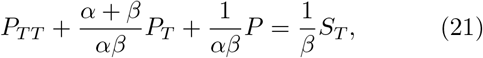

which is that of a damped harmonic oscillator, forced by the time derivative *S*_*T*_ of the signal. Recalling that *α* = |*ω*_3_| *τ*_*a*_ = 2*πτ*_*a*_*/t*_*rot*_, where *t*_*rot*_ is the rotation period, we deduce that for *τ*_*a*_*/t*_*rot*_ *<* 1 (*α <* 2*π*) there is sufficient time during a half-rotation for the adaptive response to reset itself and produce an antisymmetric response to the changing light levels, and the smaller the value of *α* the more accurately antisymmetric is the response and the more *P* mirrors the time derivative of the signal.

Moreover, the origin of the trajectories in Fig. 6(c) that head toward the light can also be understood as a consequence of the confinement of the lensed light to a small angular domain; the photoreceptor of a cell that swims in a direction *φ* significantly larger than *π/*2 will receive direct illumination during one half of the rotational cycle and will be in the dark region during the other half. This phenomenon is thus analogous to the situation described recently [21] when *Chlamydomonas* is illuminated by two nearly antiparallel sources. When the eyespot is displaced from the cell midplane, there is a range of swimming directions within which only a single source is sensed.

We have presented numerical results for the case in which a wild type cell exhibits positive phototaxis, corresponding to the choice *P* * *<* 0 [9], and lensing effects induce negative phototaxis. The symmetry of the problem is such that to describe a negatively phototactic wild type cell requires only changing the sign of *P* *, and lensing effects would then induce positive phototaxis.

In order to interpret the experimental results of Ueki, et al. [19], where cell accumulation at the edge of a quasitwo-dimensional algal suspension in a Petri dish was used as a measure of phototactic movement, we apply the twodimensional theoretical results discussed above. These imply that for a given initial position and orientation within the Petri dish, positively phototactic cells subjected to a collimated beam of light would swim away from the light asymptotically along one of two rays at ± *φ*\*, and they would intersect the edge of a petri dish at one of two spots separated by the angle 2*φ*\*. Averaging over initial positions and orientations will produce a continuous distribution of accumulation at the Petri dish edge, as seen in experiments. This smearing effect will be enhanced by at least four effects: (i) the distribution of photoreceptor positions relative to the cell equator [21], (ii) the finite size of the photoreceptor (which will smear out the caustic), and (iii) the varying photoreceptor tilts relative to the swimming direction due to a distribution of helical trajectories, and (iv) the inherent noisiness of the trajectories due to the “run-and-turn” locomotion associated with stochastic switching between periods of flagellar synchrony and asynchrony.

## IV. DISCUSSION

In this paper we have provided a quantitative theory for the experimentally observed sign reversal of phototaxis in mutants of the unicellular green alga *Chlamydomonas* that do not have the pigmented eyespot responsible for directional sensitive of the photoreceptor. Our results indicate that the sign reversal arises not from the brightness of the lensed light itself, but rather from the greater rate of change of the intensity of lensed light experienced by the photoreceptor as a cell turns. The analysis predicts that reversed-sign phototaxis is associated with swimming away from the light at a particular angle, and that a population of positively phototactic cells with randomized initial positions and orientations would display bulk movement away from the light within a broad angular distribution, as observed experimentally, but with a subset of the population moving toward the light. While existing studies [19] of the relevant phototactic mutants have not studied single-cell swimming with sufficient detail to quantify individual trajectories and test these predictions, 2D and 3D tracking methods of the kind that have been developed for studying protists [9, 10, 24] can be used.

## ACKNOWLEDGMENTS

This work was supported in part by a Trinity College Summer Studentship and a Gates Scholarship (M.Y.), Gordon and Betty Moore Foundation Grant No. 7523 (S.K.B. & R.E.G.) and Wellcome Discovery Award 307079/Z/23/Z (A.I.P. & R.E.G.).

